# Kipoi: accelerating the community exchange and reuse of predictive models for genomics

**DOI:** 10.1101/375345

**Authors:** Žiga Avsec, Roman Kreuzhuber, Johnny Israeli, Nancy Xu, Jun Cheng, Avanti Shrikumar, Abhimanyu Banerjee, Daniel S. Kim, Lara Urban, Anshul Kundaje, Oliver Stegle, Julien Gagneur

## Abstract

Advanced machine learning models applied to large-scale genomics datasets hold the promise to be major drivers for genome science. Once trained, such models can serve as a tool to probe the relationships between data modalities, including the effect of genetic variants on phenotype. However, lack of standardization and limited accessibility of trained models have hampered their impact in practice. To address this, we present Kipoi, a collaborative initiative to define standards and to foster reuse of trained models in genomics. Already, the Kipoi repository contains over 2,000 trained models that cover canonical prediction tasks in transcriptional and post-transcriptional gene regulation. The Kipoi model standard grants automated software installation and provides unified interfaces to apply and interpret models. We illustrate Kipoi through canonical use cases, including model benchmarking, transfer learning, variant effect prediction, and building new models from existing ones. By providing a unified framework to archive, share, access, use, and build on models developed by the community, Kipoi will foster the dissemination and use of machine learning models in genomics.

## Introduction

Advances in machine learning, coupled with rapidly growing volumes of molecular data, are catalyzing progresses in genomics. In particular, predictive machine learning models, which are mathematical functions trained to map input data to output values, find widespread usage including variant calling from whole genome sequencing data^1,2^, predicting CRISPR guide activity^3,4^, and predicting molecular phenotypes from the DNA sequence, including transcription factor binding, chromatin accessibility and splicing efficiency^5,6,7,8,9,10^. Once trained, such models hold the promise to allow for probing regulatory dependencies in silico, which, besides other applications, enables interpreting functional variation in personal genomes and rationalizes the design of synthetic genes.

However, despite the pivotal importance of predictive models in genomics, it is surprisingly difficult to share and exchange models effectively. In particular, there is no established standard for sharing *trained models*, in contrast to bioinformatics software and workflows, which are commonly shared through general-purpose community software platforms such as the highly successful Bioconductor project^11^, or to genomic raw data, which can be shared via data repositories such as GEO^12^, ArrayExpress^13^ and the European Nucleotide Archive^14^. Instead, trained genomics models are made available through scattered channels, including code repositories, supplementary material of articles and author-maintained web pages. The lack of a standardized framework for sharing trained models in genomics hampers their effective use, including their application to new data, and their use as building blocks to solve more complex tasks.

Repositories of trained models have helped to overcome these challenges in other fields. For example, model repositories in computer vision and natural language processing^15–17^ are routinely used for benchmarking and as a starting point to rapidly develop new models. A model repository for genomics requires additional developments in order to cover a wide range of data types of diverse genomics technologies, each of which requires specific data pre-processing strategies. A second challenge is the heterogeneity of machine learning frameworks that are currently used in the field, including Keras^18^, Tensorflow^19^, PyTorch^20^, and custom model code. Additionally, applications in genomics pose requirements on the interpretability of models, for example to understand changes in phenotype for different DNA sequence inputs. Finally, a repository of trained models for genomics needs to be easy to use and deliver robust and well-documented software to enable application by the many practitioners not expert in machine learning.

## Results

Here we present *Kipoi* (Greek for gardens, pronounced “Kípi”), a collaborative initiative to foster sharing and re-use of trained models. Already, the Kipoi repository (Fig. 1, middle) contains over 2,000 trained models that cover key predictive tasks in genomics, including the prediction of chromatin accessibility, transcription factor binding, and alternative splicing from DNA sequence. It is accessible via GitHub and the Kipoi website (https://www.kipoi.org), which provides model overviews and convenient model search functionalities. One of the core innovations of Kipoi include standardized data handling (“data-loaders”) (Fig. 1, left), which facilitates standardized data input of genomic data types across a wide range of models. Kipoi defines an application programming interface (API) (Fig. 1 right), i.e. a standard way for software components to communicate with Kipoi models that allows programmers to interchangeably use Kipoi models in their software with minimal coding effort. The Kipoi API is available in two of the most popular programing languages in bioinformatics, python and R, and from the command line, allowing any bioinformatics pipeline to integrate Kipoi models. In addition to making model predictions using established bioinformatics formats, most of the current Kipoi models (78%) can score the impact of genetic variants, and thus facilitate their functional interpretation.

**Figure 1.**
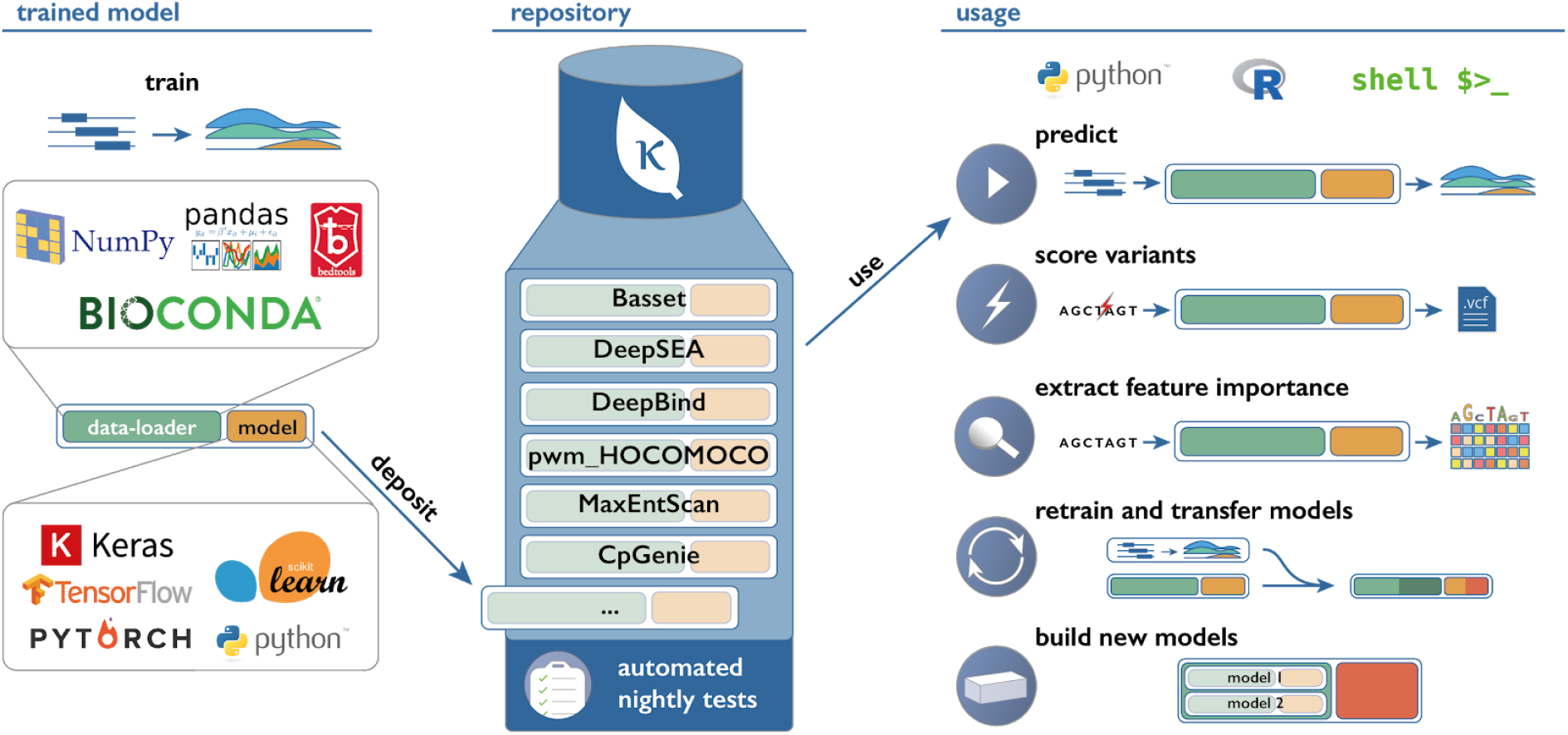
Overview of Kipoi. From left to right: At its core, Kipoi defines a programmatic standard for data-loaders and predictive models. Data-loaders translate genomics data types into numeric representation that can be used by machine learning models. Kipoi models can be implemented using a broad range of machine learning frameworks. The Kipoi repository allows community users to store and retrieve trained models together with associated data-loaders. Kipoi models are automatically versioned, nightly tested and systematically documented with examples for their use. Kipoi models can be accessed through unified interfaces using python, R, and command line to install models and all required software dependencies. Kipoi streamlines the usage of trained models to make predictions on new data, to score variants stored in standard personal genome file format, and to assess the effect of variation in the input to model predictions (feature importance score). Moreover, Kipoi models can be adapted to new tasks by retraining or by building new composite models that combine existing ones. Newly defined models can be deposited in the repository.

To support sustainability of the trained models and facilitate their dissemination, Kipoi builds on and interoperates a range of software development technologies and standards. Kipoi’s infrastructure is fully open-source: The models and the code of Kipoi itself are stored on GitHub, a code repository with issue tracking that facilitates transparent and rapid user-developer iterations. Moreover, GitHub tracks and indexes all versions of the code and models, hence facilitating the reproduction of a given analysis at any time point in the future as required for reproducible science^21^. Kipoi offers seamless installation of the models and their software dependencies independently of the programming language of the model (using Conda and pip package managers hence leveraging the Bioconda distribution^22^), addressing a major hurdle preventing the widespread sharing of trained machine learning models across the bioinformatics community. Moreover, nightly tests on all models are performed using a continuous integration service (CircleCi) to ensure model executability on test data at all times. Here, we illustrate usage of Kipoi through realistic use cases and make the code available for each of them.

### Benchmarking of alternative models predicting transcription factor binding

Practitioners are often faced with multiple predictive models for a particular task. Choosing the most appropriate model often requires a customized benchmark as the original publications describing these models typically use different datasets and provide setups favoring the published model. Access to a wide range of models through a common API facilitates such systematic comparisons. To illustrate this use case, we benchmarked five commonly used models for predicting genomic binding sites of transcription factors (Fig. 2a). These models span different modeling paradigms, including methods based on classical position weight matrices (PWM), gapped k-mer support vector machines (lsgkm-SVM^24^) and deep learning (DeepBind^5^, DeepSEA^6^ and FactorNet^7^). The models were assessed for distinguishing bound from unbound regions, where bound regions were defined as high-confidence binding events in chromatin immunoprecipitation sequencing (ChIP-seq) experiments of four transcription factors in different cell lines: CEBPB in HeLa-S3, JUND in HepG2, MAFK in K562, and NANOG in H1-hESC (Methods). The Kipoi implementations for all models except lsgkm-SVM were derived from implementations provided by the respective publications and were hence trained by the authors. The performance was assessed on chromosome 8 which was not used to train any of the considered models.

**Figure 2.**
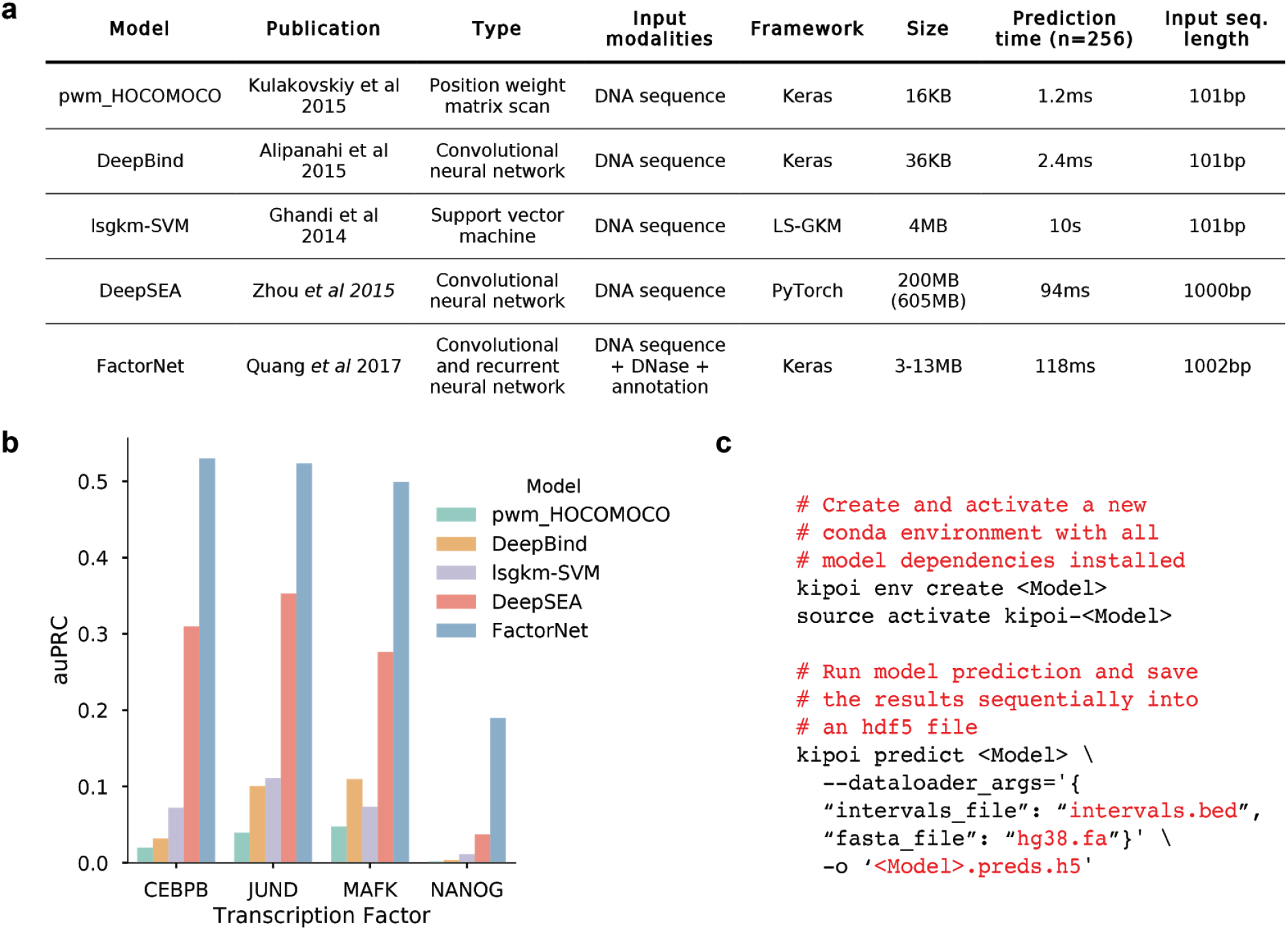
Applying and benchmarking alternative Kipoi models for transcription factor binding prediction. (a) Five models for predicting transcription factor binding that are based on alternative modeling paradigms: i) predefined position weight matrices contained in the HOCOMOCO database^23^; ii) lsgkm-SVM^24^, a support vector machine classifier; iii) the convolutional neural network DeepBind^5^; iv) the multi-task convolutional neural network DeepSEA; v) FactorNet, a multimodal deep neural network with convolutional and recurrent layers that further integrates chromatin accessibility profile and genomic annotation features. Models differ by i) the size of genomic input sequence, where DeepSEA^6^ and FactorNET^7^ consider ~1 kb sequence inputs, whereas other models are based on ~100 bp, and ii) parametrization complexity with the total size of model parameters ranging from 16kB (pwm_HOCOMOCO) to 200 Mb (DeepSEA). **(b)** Performance of the models in **a** for predicting ChIP-seq peaks of four transcription factors on held-out data (chromosome 8), quantified using the area under the precision-recall curve. More complex models yield more accurate predictions than basic models which are commonly used, **(c)** Example access to Kipoi models via the command line interface to install required software dependencies, download the model, extract and pre-process the data, and write predictions to a new file. Results as shown in **b** can be obtained for all Kipoi models using this generic command. Placeholder <Model> can be any of the models listed in **a.**

Position weight matrices generally performed poorly across all transcription factors (Fig. 2b), likely due to their inability to account for additional sequence features, such as motifs of other cooperating and competing transcription factors. More complex models (e.g. DeepSEA and FactorNet) consistently outperformed simpler ones (e.g. DeepBind and lsgkm-SVM). FactorNet yielded the most accurate predictions across all transcription factors, highlighting the importance of explicitly integrating target cell-type specific chromatin accessibility profiles with DNA sequence for predicting *in vivo* transcription factor binding (Fig. 2b). Consistent with this, we also observed that DeepSEA and FactorNet perform similarly when model evaluation is restricted to bound and unbound regions that strictly overlap accessible chromatin regions (Supp. Fig. 1, Methods).

In this example, Kipoi turned an otherwise cumbersome task into executing three simple commands (Fig 2c). The considered models are implemented using different software frameworks (Fig 2a), require different input file formats and return predictions in different formats. Additionally, installing the appropriate software dependencies for each model is difficult and time consuming without Kipoi.

### Improving predictive models of chromatin accessibility using transfer learning

Training new models can be time consuming and require large training datasets. It can be facilitated by transfer learning, i.e. by reusing models trained on one prediction task to initialize a new model for a different but related task^25^. Transfer learning typically enables more rapid training, requires less data to train and improves the predictive performance compared to models trained from scratch^26^. One class of predictive models well suited to transfer learning are deep neural networks. Deep neural networks consist of successive layers which transform input data into increasingly abstract representations. Most of the low-level abstractions, for instance edge detection for images or transcription factor motifs in genomics, turn out to be common to multiple prediction tasks. Hence, the training on a different task can be focused on the most abstract layers. Transfer learning of deep neural networks has been successfully used across multiple domains including biological imaging^27–30^, natural language processing^31^, and genomics^32^.

Here, we revisited the transfer learning example in genomics^32^ on a larger dataset of chromatin accessibility profiles for 431 biosamples (cell lines or tissues, Methods). We trained a genome-wide model predicting chromatin accessibility for 421 biosamples (tasks) while holding out 10 biosamples. For the 10 held-out biosamples, we transferred the model parameters to a new model and replaced the final layer with a randomly initialized one (Fig. 3a). One transferred single-task model was trained for each of the 10 held-out biosamples, keeping the model parameters of all layers except the last two layers fixed during re-training. Models initialized with transferred model parameters yielded improved predictive accuracy for all biosamples with 15.2% larger area under the precision-recall curve on average, compared to the same model initialized entirely with random parameters (Fig. 3b). In addition to improved performance, the training time for transferred models was substantially lower. On average, training the randomly initialized model to optimal performance required 17.3 iterations over the whole training dataset (epochs) (>1 day training time), compared to 2.8 epochs (~4 hours training time) for transferred models (Fig. 3c).

**Figure 3:**
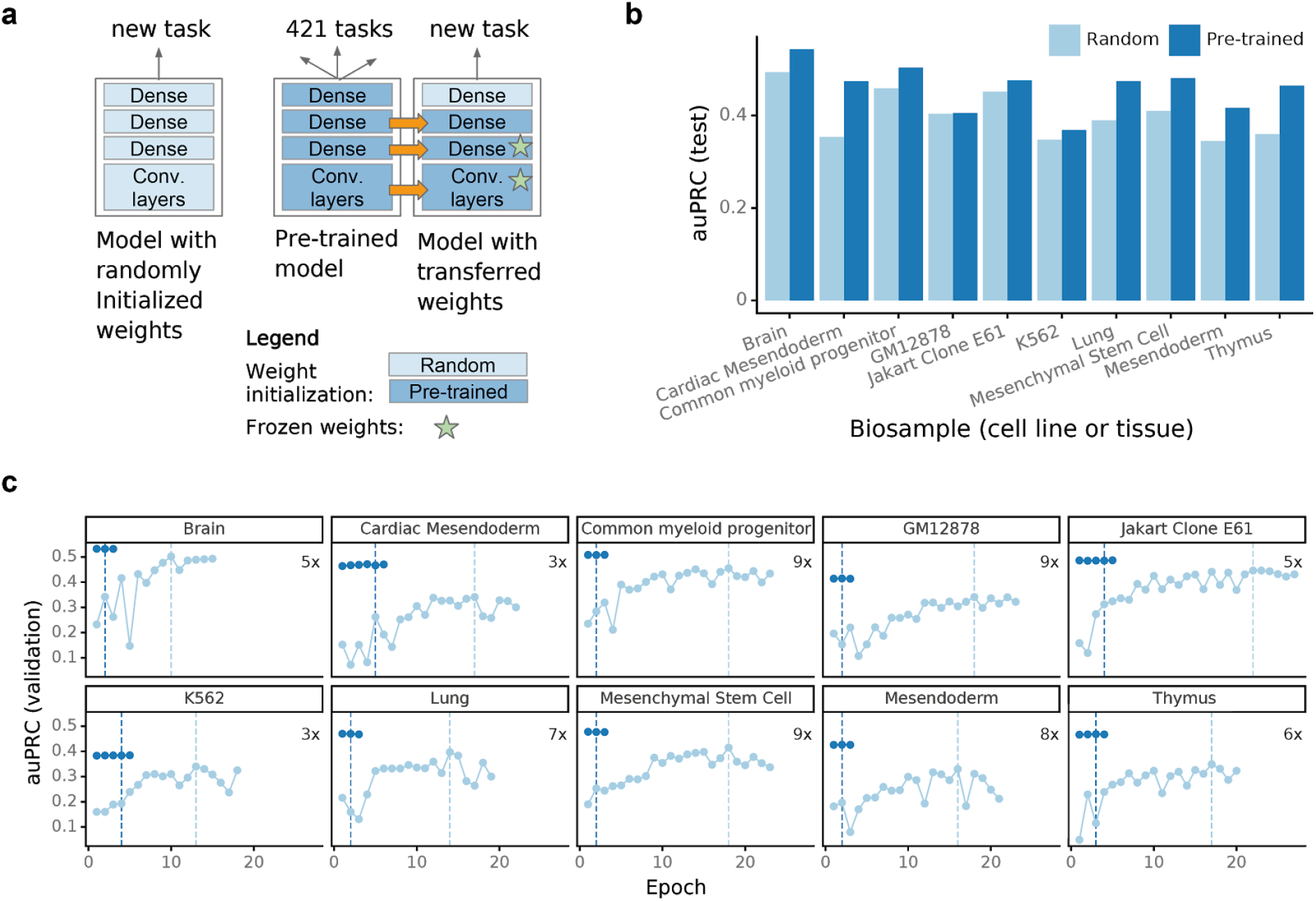
Adapting existing models to new tasks (transfer learning) **(a)** Architecture of alternative models for predicting chromatin accessibility from DNA sequence. Model parameters are either randomly initialized (left) or transferred from an existing neural network pre-trained on 421 other biosamples (cell lines or tissues, right), **(b)** Prediction accuracy measured using the area under the precision-recall curve, comparing randomly initialized (light blue) versus pre-trained (dark blue) models. Shown is the performance on held-out test data (chromosomes 1, 8 and 21) for 10 biosamples that were not used during pre-training, **(c)** Training curves, showing the area under the precision-recall curve on the validation data (chromosome 9) as a function of the training epoch. The dashed vertical line denotes the training epoch at which the model training is completed. Pre-trained models require fewer training epochs than randomly initialized models and they achieve more accurate predictions.

Kipoi promotes transfer learning in three ways. First, it provides access to a comprehensive collection of state-of-the-art models in genomics. Transfer learning works well if the tackled task is similar to the original task of the pre-trained model^25^. Kipoi allows users to quickly browse models by name, tag or framework and hence find the model candidate closest to their task at hand. Second, each model is easily installable and comes packaged with a tested data-loader. Most of the data-loaders can be directly used to re-train models. Third, for neural network models, Kipoi offers a command to return and store the activation of a desired intermediate layer rather than the final, prediction layer. The transferred model can take those activations as input features instead of the original input. Since the intermediate layer can serve as a good feature extractor, this procedure can speed up the training process by multiple orders of magnitude without reducing performance. For the transfer learning example in Fig. 3, model training took only 3 minutes with the pre-computed values on a single graphical card (NVIDIA TITAN X). Altogether, leveraging pre-trained models, in particular deep models that have been trained on large datasets with a substantial investment in compute time, allow researchers to train more accurate models on smaller datasets while saving time and compute costs.

### Predicting the molecular effects of genetic variants using interpretation plugins

One important application of trained models in genomics, with translational relevance in human genetics and cancer research, is to predict the effects of genetic variants on molecular phenotypes’^6,33^. Individually, variant effect prediction has been implemented by a subset of published sequence-based predictive models such as DeepBind^5^, DeepSEA^6^, and CpGenie^33^. In Kipoi this is generalized and implemented as a plugin that allows annotating variants obtained from the variant call format (VCF) files using any DNA sequence based model. The variant effect prediction plugin performs in-silico mutagenesis by contrasting model predictions for the reference allele and for the alternative allele (Fig. 4a). If the model can be applied across the entire genome, such as chromatin accessibility models, sequences centered on the queried variants are extracted (top row, Fig. 4b). If instead the model can only be applied to regions anchored at specific genomic locations, such as splicing models at intron-exons junctions, only sequences extracted from valid regions that overlap with the variants of interest are used (bottom row, Fig. 4b). A uniform handling of these two scenarios using a single command (Fig. 4c) greatly simplifies their application. Altogether, the variant effect prediction plugin allows integrating a broad range of regulatory genomics predictive models into personal genome annotation pipelines and is trivially extended with newly added models.

**Figure 4:**
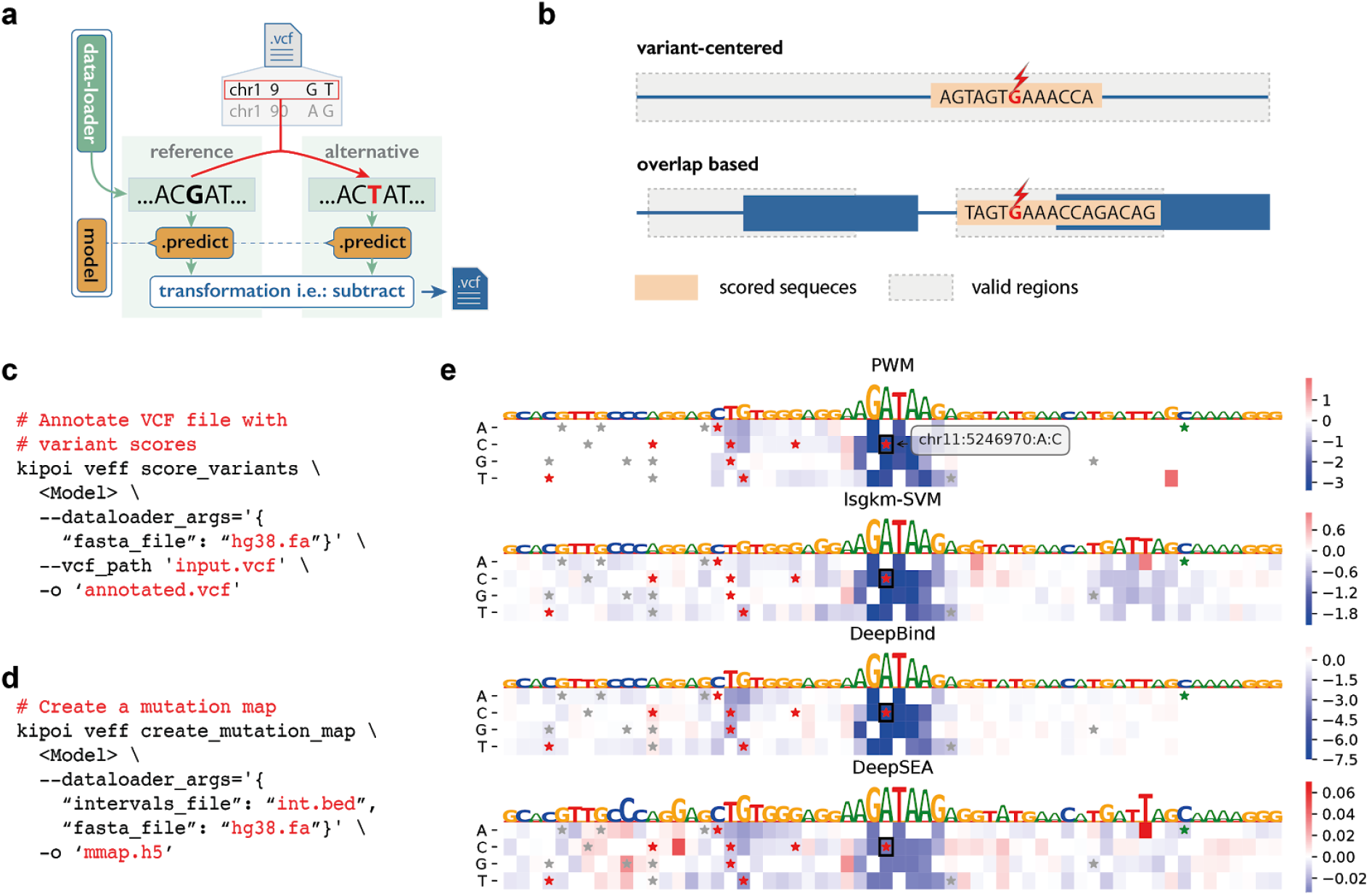
Variant effect prediction and feature importance scores. **(a)** Schema of variant effect prediction using in-silico mutagenesis. Model predictions calculated for the reference allele and the alternative allele are contrasted and written into an annotated copy of the input variant call format file (VCF). **(b)** Kipoi uniformly supports variant effect prediction for models that can make predictions anywhere in the genome (top) and also for models that can make predictions only on predefined regions such as exon boundaries (bottom), **(c)** Generic command for variant effect prediction, **(d)** Generic command to compute the importance scores using in-silico mutagenesis **(e)** Feature importance scores visualized as a mutation map (heatmap, blue negative effect, red positive effect) for variant rs35703285 and the predicted GATA2 binding difference between alleles for 4 different models. The black boxes in the mutation maps highlight the position and the alternative allele of the respective variant. Additionally, stars highlight variants annotated in the human variant database ClinVar with red: (likely) pathogenic, green: likely benign, grey: uncertain or conflicting significance, other.

To inspect genomic regions containing the variant in higher detail, variant effect predictions for all possible single nucleotide variants in the sequence can be computed using a single command (Fig. 4d) and visualized as a mutation map (Fig. 4e). This helps to assess the predicted impact of the variant of interest in the context of other possible variants in the genomic region and may help pinpoint the affected cis-regulatory elements. For example, the mutation maps for transcription factor binding sites of GATA2 show that the first four models from Fig. 2 agree on the effect of the variant rs35703285. Interestingly, the three most complex models (lsgkmSVM, DeepBind, and DeepSEA) predict effects of similar strength further away from the core motifs. This reflects that they can model more complex regulatory structure than the sole core motif captured by the position weight matrix approach. Variant rs35703285 has been classified as pathogenic in the ClinVar dataset and is linked to beta Thalassemia (MedGen:C0005283), a disease that reduces synthesis of the hemoglobin subunit beta (hemoglobin beta chain) that results in microcytic hypochromic anemia^34^. The mutation map illustrates that similar loss of GATA2 binding can be expected from other variants in the region.

In addition to in-silico mutagenesis, which only applies to sequences, Kipoi provides a plugin that can evaluate the influence for any type of input on model prediction by implementing various feature importance algorithms, including saliency maps^35^ and DeepLift^36^. These feature importance algorithms offer an additional perspective and are often much faster to compute than in-silico mutagenesis.

### Predicting pathogenic splice variants by combining models

State of the art models performing variant effect prediction frequently combine scores from multiple models. The advantage is two-fold. First, combined scores can cover multiple biological processes. Second, combined scores are more robust, because they average out conflicting predictions of individual models. Combining models or scores can be easily done in Kipoi by leveraging the standardization and modularity of models in combination with the variant effect prediction plugin introduced above. As a proof-of-concept, we used Kipoi to define a pathogenicity score of variants located near splice sites by integrating four predictive models covering complementary aspects of splicing (Fig 5a) into a single composite model.

Defect in splicing is one of the most frequent cause of genetic disease (López-Bigas et al., 2005). In the first step of splicing, the donor site is attacked by an intronic adenosine to form a branchpoint. In the second step, the acceptor site is cleaved and spliced (i.e. joined) to the 3’ end of the donor site. To cover variants possibly affecting splicing through different mechanisms, we considered four complementary models trained on different types of data. These models were i,ii) 5’ and 3’ MaxEntScan^8^, a probabilistic model scoring donor and acceptor site regions that was trained on splice sites with cDNA support, iii) HAL^9^, a k-mer based linear regression model scoring donor sites that was trained on a massively parallel reporter assay in which hundreds of thousands of random sequences probed the donor site sequence space^9^, and iv) Labranchor, a deep-learning model scoring the region upstream of the acceptor site for possible branchpoint locations that was trained from experimentally mapped branchpoints^37^.

While MaxEntScan can be easily applied to score genetic variants provided in VCF files through ENSEMBL’s variant effect predictor plugin^38^, HAL and Labranchor do not offer this functionality out-of-the-box. Using Kipoi’s API, the variant effect prediction is standardized for all these models (Fig. 5a). We built a new Kipoi model, KipoiSplice4, which is a logistic regression model based on variant effect predictions of these four Kipoi models and phylogenetic conservation scores (Methods, Fig 5a). This combined model was trained on two different datasets of splice variants classified either as pathogenic or benign (dbscSNV and ClinVar, Methods).

**Figure 5:**
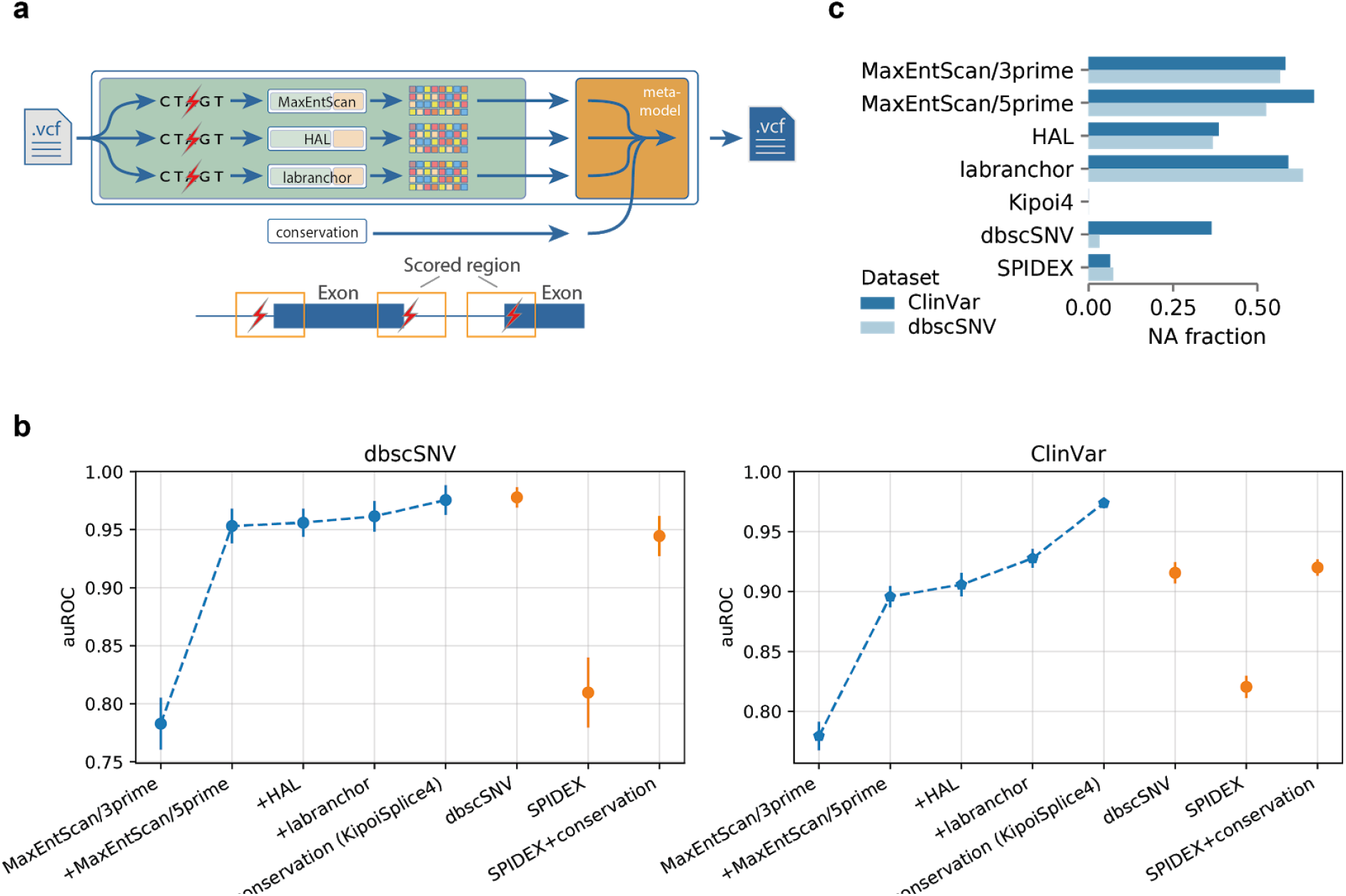
Composite models using Kipoi for improved pathogenic splice variant scoring. Illustration of composite modelling for mRNA splicing. A model trained to distinguish pathogenic from benign splicing region variants is easily constructed by combining Kipoi models for complementary aspects of splicing regulation (MaxEntScan 3’ models acceptor site, MaxEntScan 5’ and HAL model donor sites, labranchor models the branchpoint) and phylogenetic conservation. These variant scores are combined by logistic regression to predict the variant pathogenicity (orange box), **(b)** Different versions of the ensemble model were trained and evaluated in 10-fold cross-validation for the dbscSNV and ClinVar datasets (Methods). The four leftmost models are incrementally added to the composite model in chronological order of their publication: the leftmost point only uses information from the MaxEntScan/3prime model, while '+conservation (KipoiSplice4)' uses all four models and phylogenetic conservation. These performances were compared to a logistic regression model using state-of-the-art splicing variant effect predictors (SPIDEX, SPIDEX+conservation, dbscSNV). KipoiSplice4 achieves state-of-the-art performance on the dbscSNV dataset and outperforms alternative models on ClinVar which contains a broader range of variants **(c)** Fraction of unscored variants for different models in the dbscSNV and ClinVar datasets.

To illustrate the benefit of integrating multiple models, we incrementally added the four splicing models in the chronological order of model publication. With an increasing number of models, the performance increased in both, dbscSNV and ClinVar datasets (Fig. 5b, four left-most methods). Next, we evaluated the model performance against two state-of-the-art splicing scores: another integrative approach that predicts pathogenic splicing-affecting variants dbscSNV^39^ and SPIDEX^40^. For a fair comparison, we furthermore trained a score combining SPIDEX and phylogenetic conservation on each dataset, which reached the same performance as the dbscSNV model on ClinVar. While the performance of KipoiSplice4 is similar to dbscSNV for the dbscSNV dataset, KipoiSplice4 outperforms all other methods on the ClinVar dataset. One reason for the better performance of KipoiSplice4 is that it scores more variants (Fig. 5c). Neither SPIDEX nor dbscSNV explicitly model the splicing branchpoint, while KipoiSplice4 does so using labranchor.

By wrapping the individual models into a data-loader, we made the ensemble model KipoiSplice4 available in Kipoi. KipoiSplice4 can hence be executed on demand to *de novo* predict effects of variants in splice sites. This is not possible with other state-of-the-art splicing models which are published as pre-computed databases, such as dbscSNV. While SPIDEX offers a web interface to generate variant scores, it can currently only do so for 40 variants at a time. Altogether, by wrapping existing splice-models into Kipoi, and thereby leveraging the out-of-the-box variant effect prediction, we developed a state-of-the-art model for scoring the pathogenicity of splicing variants. Additionally, with new splicing models and more extensive training datasets of better quality being published, the ensemble model can be easily and transparently improved.

## Discussion

We have developed a repository and programmatic standard for sharing and re-use of trained models in genomics, thereby addressing an unmet need. By providing a unified interface to models, automated installation, and nightly tests, Kipoi streamlines the application of trained models, overcomes the technical hurdles of their deployment, improves their dissemination, and ultimately facilitates reproducible research. The use cases presented demonstrate that Kipoi greatly facilitates the execution and comparison of alternative models for the same task, standardizes their use to functionally interpret genetic variants, and facilitates the development of new models based on existing ones, either by means of transfer learning or by model combination.

The dissemination and sharing of trained models has major advantages compared to sharing pre-computed predictions or to sharing code for users to train models from scratch. In particular, pre-computed predictions cannot be extended to new or different input data. Moreover, the generation of extensive sets of pre-computing results for a wide range of potentially relevant input values can be prohibitive in terms of compute time and storage. For example, storing variant effect predictions is technically impossible even for relatively short (<10bp) indels for combinatorial reasons. On the other hand, re-training models from scratch is frequently non-trivial, requires access to potentially very large training dataset, and can require large computational resources. Trained machine learning models can be regarded as functions encoding data distributions^41^. Hence, it is maybe not surprising that a demand for repositories dedicated to trained models arises in the era of big data, where they fill a gap between code repositories and data repositories.

At the core of our contribution is an application programmatic interface (API), a unified way for software components to interact with any of these models. APIs provide modularity to software design, help to reduce code redundancy and allow developers to focus only on the most relevant tasks. We demonstrated the utility of the API, which provides a generic approach to carry out variant effect predictions, and to derivefeature importance scores for a wide range of models. These examples are important downstream functionalities which are not typically provided by software implementations of models as provided by authors, or they may be implemented using diverse and inconsistent paradigms and interfaces. We foresee a range of future plugins that are of general use for different models. While most of the models in Kipoi currently predict molecular phenotypes from DNA sequence, the design of Kipoi is agnostic to input or output data types. Additionally, the API can be used with multiple model repositories, both public and private, simultaneously. Hence, the genericity of Kipoi makes it attractive for applications beyond the domain of genomics.

While complying to a programmatic standard can constrain contributors and provide some initial overhead to adapt legacy software, the long-term community benefits from the standardization will outweigh short-term investments. The open software project Bioconductor and the data repository GEO are canonical examples of the expected gains. These frameworks achieve a suitable compromise between rigidly enforced structure and no structure. With this in mind, we have designed Kipoi’s API to rigorously specify specific aspects such as providing example files to test model executability, while leaving other choices, such as the machine learning modelling framework, opento developers. We anticipate that community usage will help to develop good practices and find a reasonable balance between standardization and flexibility.

An exciting next step would be to set up open challenges for key predictive tasks in genomics with open challenge platforms like DREAM^42^ or CAGI^43^, and make the best models available in Kipoi. This would simplify and modularize the development of predictive models into three steps: (1) designing training and evaluation datasets (challenge organizers), (2) training the best model (challenge competitors) and (3) making the model easily available for others to use (repository of trained models). Such modularization would lower the entry barrier for newcomers as well as machine learning practitioners lacking domain expertise. Moreover, as models and training datasets continue to evolve, such best-in-class models could be continuously updated and made immediately available to all. Kipoi provides important elements to this end: a standardization for data loading and model execution, nightly tests, and a central repository.

A repository of interoperable models opens the possibility of building composite models that capture how genetic variation propagates through successive biological processes. Such a sequential, modular modeling offers multiple advantages. First, end-to-end fitting of a complex trait such as a cellular behavior or the expression level of a gene can be too difficult because the amount of data is too scarce compared to the complexity of the phenomena. In contrast, today’s high-throughput technologies focusing on a specific sub-process offer more data at higher accuracy. For example, massively parallel reporter assays allow performing saturated screens in which nearly the complete combinatorial sequence space can be probed for the selected molecular processes. Hence accurate models may be obtained for these elementary tasks and serve as building blocks for modeling more complex tasks. Second, modularity is a hallmark of biological processes as the same proteins are often involved in multiple processes. We therefore anticipate fruitful cross-talks between modelers sharing individual components useful for different modeling tasks. Third, such approach would lead to models that are interpretable in terms of simpler biological processes as opposed to black box predictors. Whether and how predictive models of elementary steps can be sequentially combined and fitted together to model multiple higher order biological processes of increasing complexity is an exciting research direction. Altogether, we foresee Kipoi as a catalyst in the endeavour to model complex phenotypes from genotype.

## Methods

### Kipoi infrastructure

#### Model source

Kipoi’s main model repository (“model source”) is hosted as a git repository at https://github.com/kipoi/models. Each folder containing the following files is considered to be a single model:

- model.yaml - model description in the YAML format
- model_files/ - directory with files required by the model (like model weights)
- model.py - (optional) model implementation as a python class
- example_files/ - directory with small files used to test the model
- dataloader.py - data-loader implementation (could be implemented by other models)
- dataloader.yaml - data-loader description (could be implemented by other models)
- dataloader_files/ - (optional) directory with files required by the data-loader

The ‘model.yaml’ and ‘dataloader.yaml’ each specify (i) general information like author name, publication link or description, (ii) required software dependencies, (iii) input-output data types, and (iv) additional information required by plugins like variant effect prediction, ‘model_files’, ‘dataloader_files’, ‘model.py’ and ‘dataloader.py’ contain the code and parameters necessary to execute the model, ‘example_files’ contains small input files to test the execution of model prediction.

Folders whose name ends with ‘_files’ are tracked by Git Large File Storage (LFS, https://git-lfs.aithub.com/). In addition to Kipoi’s default model source, the user can host and seamlessly use their own, private or public, model source. Model sources are specified in Kipoi’s config file and are treated completely equivalently to the default model source.

#### Depositing and testing models

New models or updates to existing models are submitted as pull requests to the Kipoi model repository https://github.com/kipoi/models. For each pull request, the added or updated models are automatically tested using the CircleCI continuous integration service. Additionally, all models from the master branch of the repository are automatically tested every day. For each tested model, a new Conda environment with all the required dependencies will get installed, and model prediction will get executed for the example files. In order to pass the tests no errors or warnings can be raised, and the arrays returned by both, the data-loader and the model, have to be consistent with the description in their yaml files.

#### The API

Kipoi’s API is implemented as a python package supporting python 2.7 and python>=3.5. The package is directly installable from PyPI and Bioconda^22^. It provides a command line interface exposed through the ‘kipoi’ command. Using Kipoi from the R programing language is enabled by using the ‘reticulate’ R package. The API provides functionality necessary to manage model and data-loader dependencies, it provides generic methods for executing model predictions and gradient calculations (where available). To enable generic definition of interfaces Kipoi defines two main classes: ‘Model’ and ‘Dataloader’.

#### Model

Model is a class implementing the method `predict_on_batch (x)`. Argument x can be a single numpy array, a list of numpy arrays or a dictionary of numpy arrays. In its current version, Kipoi wraps models implemented in Keras, Tensorflow, PyTorch and Scikit-learn. For models developed in one of these frameworks, the contributor can directly provide the serialized model. A user can also deposit a custom model by implementing the Model class and hence make use of arbitrary python code or even command-line calls. For models implemented in deep learning frameworks (Keras, Tensorflow, PyTorch), the model class additionally provides two methods: `predict activation on batch (x, layer, pre_noniinearity= False)`, which returns the feature activation map of an intermediary layer (useful for transfer learning) and `input_grad (x, fiiter_idx=None, avg_func=None, wrt_iayer=None, …)`, which returns the gradient of the input with respect to model’s predictions (useful for feature importance scores). Support for additional machine learning frameworks can be easily added.

#### Data-loader

The aim of the data-loader is to generate batches of data consumable by the model. It encapsulates the loading of data from input files and its pre-processing. The data-loader has to return a dictionary with three keys: inputs, targets (optional), metadata (optional). Value of the ‘inputs’ key is directly passed on to the model input. ‘targets’ provide labels useful for training or benchmarking. ‘metadata’ optionally provide additional information about the data samples (like sample identifier or genomic ranges of the extracted genome sequence).

To implement a data-loader, the contributor can either write a python function, generator, iterator or a Dataset class (http://kipoi.org/docs/contributing/04Writingdataloader.pv/). Regardless of how the data-loader is implemented, the user will have direct access to the following methods: `batch_iter` returning batches of data stored as a dictionary with inputs, targets and metadata keys, `batch_train_iter` returning batches of data indefinitely as a tuple of inputs and targets (directly useful with the Keras’ fit_generator), `batch_predict_iter` returning batches of inputs and `ioad_aii` returning the whole dataset. Parallel data-loading is by default enabled for data-loaders written as a ‘Dataset’ by using the DataLoader class originally implemented in PyTorch.

#### Variant effect prediction and model interpretation plugins

Additional domain-specific functionality of models can be implemented in the form of additional python packages - Kipoi plugins. We implemented two plugins: variant effect prediction (https://github.com/kipoi/kipoi-veff) based on in-silico mutagenesis and model interpretation using feature importance scores (https://github.com/kipoi/kipoi-veff).

#### Dependency installation

Model and the data-loader can specify dependencies installable either by the Conda package manager (https://conda.io) or the ‘pip’ (https://pypi.org/proiect/pip) package manager. Thanks to the open source efforts like conda-forge (https://conda-forae.org/) or Bioconda (https://biocond.github.io/), the Conda package manager covers a large set of dependencies including all major bioinformatics packages. Since Conda is a package manager for any programming language, not just python, it is easy to integrate models that are implemented in another language and only expose a command-line interface. One such example is lsgkm-SVM which is precompiled and distributed through the bioconda channel. One example Kipoi model built on top of lsgkm-SVM is lsgkm-SVM/Tfbs/Ctcf/K562/Uw_Std. The main strength of Conda is that it can create virtual environments. This allows the user to create multiple environments for different models. For the definition of computation pipelines we recommend using Kipoi’s command line interface and virtual environment creation with Snakemake^44^. This allows to calculate model predictions for different models using a single generic snakemake rule while the model predictions get executed in isolated environments.

### Models

#### pwm_HOCOMOCO

Position weight matrices (PWM) for all 600 human transcription factors in HOCOMOCO v10 were downloaded from http://hocomoco10.autosome.ru/final_bundle/HUMAN/mono/HOCOMOCOv10_pcms_HUMAN_mono.txt and transformed to position specific scoring matrices (PSSM) using the pseudocount probability of 0.001. Scanning the DNA sequences using the PSSM matrix is implemented as a Keras model consisting of a single convolutional layer with one filter whose weights are set to the PSSM, followed by global max pooling. The model operates on one-hot-encoded DNA sequence.

#### DeepBind

Original weights and architecture were obtained from supplementary material of the original publication^5^ and were converted to Keras 2.0 models (code: https://github.com/kundajelab/DeepBindToKeras).

#### DeepSEA

The DeepSEA model was converted from the original Torch7 model^6^ to a PyTorch model using a modified version of the script https://github.com/clcarwin/convert_torch_to_pytorch. Since prediction of model tasks and variant effect prediction use different handling of reverse-complement sequences there are two models in the Kipoi model zoo dedicated to the two different used cases in order to replicate results from the original model exactly. Implementations of reverse-complement handling were taken from .lua files provided in the software package in the publication^6^. Predictions of the models and variant effects produced by the models in the Kipoi repository match the predictions produced by the website http://deepsea.princeton.edu/job/analysis/create/.

#### FactorNet

FactorNet models were obtained from https://github.com/uci-cbcl/FactorNet. In addition to the models available in the github repository, Daniel Quang kindly provided the trained models for CEBPB and MAFK, which were part of the internal evaluation round in the ENCODE-DREAM *in vivo* transcription factor binding prediction challenge (http://svnapse.org/encode). The models were converted to Keras 2.0 supporting the tensorflow backend.

#### MaxEntScan

We used MaxEntScan implemented in the maxentpy package (https://github.com/kepbod/maxentpy) provided through the Bioconda channel. We implemented a data-loader that takes the reference genome FASTA file and the genome annotation GTF file as input and returns sequences of all regions [-3nt,5nt] w.r.t. the annotated 5’ splice sites for the 5’ model and sequences of all regions [-3nt,20nt] w.r.t. the annotated 3’ splice sites for the 3’ model.

#### HAL

The HAL model was adapted from https://github.com/Alex-Rosenberg/cell-2015 by implementing the identical 5’ splice-site scoring function into a Kipoi’s model class with the ‘predict_on_batch’ function. Model weights were obtained from the same repository and directly applied. We implemented a data-loader that takes the reference genome FASTA file and the genome annotation GTF file as inputs and returns k-mer counts of sequences from all regions [-80nt, 80nt] w.r.t. the annotated 5’ splice sites.

#### Labranchor

The Labranchor model was obtained from https://github.com/jpaggi/labranchor. The Keras model implementation provided by the authors could directly be used for the Kipoi model. We implemented a data-loader that takes the reference genome FASTA file and the genome annotation GTF file as inputs and returns one-hot-encoded sequences of all regions [-70, 0] nt relative to the annotated 3’ splice sites.

### Benchmarking transcription factor binding prediction models

The complete Snakefile for the analysis described in this section is available at https://github.com/kipoi/manuscript/blob/master/src/tf-bindina/Snakefile.

#### Data and prediction command

The test set for transcription factor binding models was generated using 101 bp contiguous intervals throughout chromosome 8 in the human genome assembly hg19. Each interval was labeled based on majority overlap with transcription factor ChIP-seq high-confidence peaks (IDR<0.05) from the ENCODE-DREAM in-vivo transcription factor binding site prediction challenge (http://svnapse.org/encode). Intervals in the hg19 blacklist regions (https://www.encodeproject.org/annotations/ENCSR636HFF were removed. CEBPB was evaluated in the HeLa-S3 (ENCFF002CSA), JUND in HepG2 (ENCSR000EEI), MAFK in K562 (ENCFF812QPN) and NANOG in H1-hESC (ENCFF379EPK) cell type. The additional files required by FactorNet (like the DNase accessibility track) were obtained from the URLs listed in https://github.com/uci-cbcl/FactorNet/tree/master/data#bigwig-files. For models that require sequence lengths of more than 101 bp, we increased the size of labeled intervals. For example, to provide 1002 bp intervals for FactorNet, we subtracted 450 bp from start coordinates and added 451 bp to end coordinates. All model predictions were obtained by running the ‘kipoi predict’ command in the individual conda environment for each model.

## Accessible-only regions (Supp. Figure 1)

In addition to chromosome-wide evaluation, the auPRC was computed only for regions overlapping DNase-seq signal peak regions in the corresponding cell-type by more than 50%. DNase-seq peaks were obtained from the relaxed peaks provided by the ENCODE-DREAM *in-vivo* transcription factor binding challenge (http://svnapse.org/encode).

### Isgkm-SVM training

lsgkm-SVM from Bioconda (bioconda7ls-gkm=0.0.1) was used for model training and prediction. The model was retrained on ENCODE datasets using files downloaded from the same source as mentioned in the publication^45^. Preprocessing of training was performed using the gkmSVM R-package using default parameters ‘genNullSeqs(…, nMaxTrials=20, xfold=1, genomeVersion-hg19’,..)‘. For training the 322 datasets with the most peaks were chosen, similar to the lsgkm-SVM publication. Training was performed with the parameters ‘gkmtrain -I 11 -d 3 -c 1 -T 16 -m 5120 -v 3’. For the final model chromosome 8 and 9 were held out from training to enable model benchmarking comparable with the DeepSEA models. Trained models reached area under the receiver operating curve (auROC) similar to the original publication^45^:

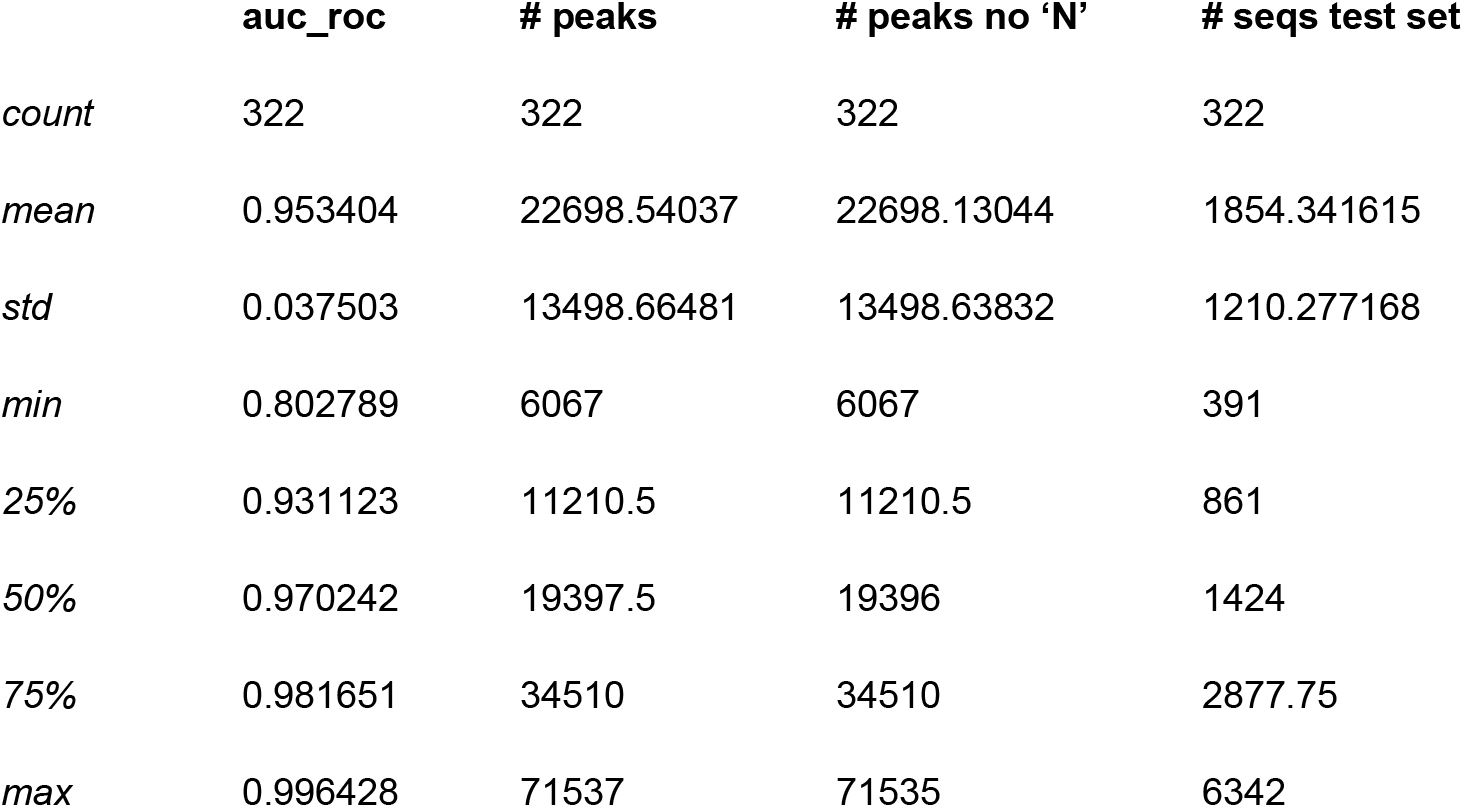

### Transfer learning

#### Peak File Acquisition

We downloaded DNase files for 431 biosamples (cell lines or tissues) from Roadmap (http://www.ncbi.nlm.nih.aov/aeo/roadmap/epiaenomics/) and ENCODE (https://www.encodeproiect.org/). and processed them separately to obtain a final dataset of binary labels (0/1) indicating chromatin accessibility in each interval of the combined accessibility region for each biosample. We processed the raw data as follows: The fastq files were aligned with BWA aln (v0.7.10), where all datasets were treated as single-end. Dynamic read trimming was set to 5, the seed length was 32, and 2 mismatches maximum were allowed in mapping. After mapping, reads were filtered to remove unmapped reads and mates, non-primary alignments, reads failing platform/vendor quality checks, and PCR/optical duplicates (-F 1804). Low quality reads (MAPQ < 30) were also removed. Duplicates were marked with Picard MarkDuplicates and removed. The final filtered file was converted to tagAlign format (BED 3+3) using bedtools’ ‘bamtobed’. Cross-correlation scores were obtained for each file using phantompeakqualtools (v1.1).

All files with a cross-correlation quality tag below 0 were discarded. For the ENCODE data generated from the Stam Lab protocol, the datasets were trimmed to 36 bp and technical replicates were combined. After removing mitochondrial and ambiguously mapped reads, the reads were randomly subsampled to a total of 50 million reads per sample. For the ENCODE data generated from the Crawford Lab protocol, the same procedure as above was performed, except reads were trimmed to 20 bp due to the different library generation protocol. For the Roadmap data, which was all generated by the Stam Lab protocol, the same procedure as above was performed with trimming to 36 bp. Reads from multiple files were combined and subsampled to 50 million reads in case the total number of reads was more than 50 million.

These trimmed, filtered, subsampled tagAlign files were then used to generate signal tracks and call peaks. Signal tracks and peaks were called with a loose threshold (p < 0.01) with MACS2 to generate bigwig files (fold enrichment and p-value) and Narrow Peak files, respectively. To obtain final peak sets, we performed pseudoreplicate subsampling on the pooled reads across all replicates (taking all reads from the final tagAligns and splitting in half by random assignment to two replicates) and running IDR (v2.0.3) with a p-value threshold of <0.1 to get a consensus region set for each DNase experiment.

#### Data Preprocessing

We divided the genome into intervals of width 1000 bp using a stride of 200 bp. For each interval, we use the hg19 reference genome to extract the DNA sequence and assign a binary label of 0 (negative) or 1 (positive) for each of the 431 biosamples if the central 200 bp of the interval overlapped at least 50% of the accessibility IDR peak or if the accessibility I DR peak overlapped at least 50% of the the central 200 bp of the interval. This resulted in the 16,551,625 intervals and 431 binary labels per interval for each of the biosamples. We use data from chromosomes 1, 8, and 21 for testing, data from chromosome 9 for validation, and the remaining data for training the models.

We selected 10 biosamples to benchmark our transfer learning procedure by performing hierarchical clustering and randomly selecting one biosample from each of the 10 clusters. Selected biosamples were: common myeloid progenitor, GM12878, Jurkat clone E61, K562, mesendoderm, mesenchymal stem cell, cardiac mesoderm, thymus, lung, and brain.

#### Model Architecture

We trained 3 types of models predicting chromatin accessibility given DNA sequence: one multi-task model with randomly initialized weights predicting accessibility for 421 cell-types, and two types of single-task models trained on the remaining 10 cell-types: a model with randomly initialized weights and a model with weights transferred from the multi-task model. All models were convolutional neural networks (CNN) with the BASSET^32^ architecture and were implemented in Keras version 1.2 using tensorflow-gpu version 1.0.0 backend.

#### Transfer Learning

We used the trained multi-task model and transferred the weights from all but the final classification layer to the transferred single-task architecture. We froze the weights of all layers but the final two, and replaced the final classification layer with a layer outputting a single prediction, instead of 421.

#### Model Training and Evaluation

Randomly initialized models were trained using a categorical or binary cross-entropy loss, batch size of 256, epoch size of 2,500,000 and the ADAM optimizer^46^ with a learning rate of 0. 0003. Pre-trained models were fine-tuned using a batch size of 128, no restrictions on the epoch size and the default learning rate (0.001) using the ADAM optimizer. Early stopping monitoring auPRC on the validation set was used with patience of 4 epochs for models with randomly initialized weights and monitoring the validation cross-entropy loss with patience of 1 epoch for models with transferred weights. Single-cell model and transferred single-cell model were evaluated on the same test set for a given biosample (chromosomes 1, 8 and 21). For example in the GM12878 biosample, the test set contains 135,630 positives and 2,330,052 negatives, and the validation set contains 35,526 positives and 647,116 negatives.

### Predicting the molecular effects of genetic variants using interpretation plugins

The presented variants were selected from the ClinVar release from April 2018. The selection involved performing variant effect prediction for all variants in the DeepSEA model and selecting the variant with the strongest negative predicted effect in GATA2 model outputs respectively. Mutation maps centered on those two variants were generated using the mutation map commands displayed in Fig. 4d and implemented in the kipoi-veff plugin.

### Predicting pathogenic splice variants by combining models

#### Data: ClinVar

The ClinVar release from April 2018 based on the reference genome GRCh37 was used (ftp://ftp.ncbi.nlm.nih.gov/pub/clinvar/vcf_GRCh37/clinvar_20180429.vcf.gz). Only variants in the range [−40nt, 10nt] around the splicing acceptor or variants in the range [-10, 10] nt around the splice donor of a protein coding gene (ENSEMBL GRCh37 v75 annotation) were used. The positive set comprises of variants classified as “Pathogenic” (6,310 variants) and the negative set comprises of variants classified as “Benign” (4,405 variants). Variants causing a premature stop codon were discarded. Per-variant pathogenicity/conservation scores (‘CADD_raw’, ‘CADD_phred’, ‘phyloP46way_placental’, ‘phyloP46way_primate’) and the dbscSNV score were obtained by VEP^38^. Spidex scores were obtained from ANNOVAR (http://www.openbioinformatics.org/annovar/spidex_download_form.php).

#### Features

##### Kipoi features

For specific Kipoi models the following features were produced:

- MaxEntScan/3prime, MaxEntScan/5prime, HAL

- <model>_ref: Model prediction for the reference allele
- <model>_alt: Model prediction for the alternative allele

- Labranchor
- labranchor_logit_ref: (optional) Model prediction on the for the reference allele on the logit scale
- labranchor_logit_alt: (optional) Model prediction on the for the alternative allele on the logit scale

- All models
<model>_is_na: 1 if model prediction is unavailable for the variant and 0 otherwise

These features were obtained by running the`kipoi veff score_variants` command on the variants table with`-s logit_ref logit_ait ref alt logit dif f` formatted as a vcf file and then parsing the returned vcf files using `kipoi veff.parsers. KipoiVCFParser`.

##### dbscSNV features

- dbscSNV_rf_score’ - dbscSNV random forest score obtained with VEP
- ‘dbscSNV_rf_score_isna −1 if dbscSNV_rf_score’ is unavailable for the variant and 0 otherwise

##### SPIDEX features

- dpsi_max_tissue’, the maximum mutation-induced change in percentage-spliced in (PSI) across 16 tissue
- ‘dpsi_max_tissue_isna’, 1 if dpsi_max_tissue’ is unavailable for the variant and 0 otherwise
- ‘dpsi_zscore’, z-score transformed dpsi_max_tissue
- ‘dpsi_zscore_isna, 1 if ‘dpsi_zscore’ is unavailable for the variant and 0 otherwise

##### Conservation features

All obtained using VEP

- CADD_raw, Combined Annotation-Dependent Depletion score as described in ^47^
- CADD_phred, CADD phred-like rank score based on whole genome CADD raw scores
- phyloP46way_placental, phyloP (phylogenetic p-values) conservation score based on the multiple alignments of 33 placental mammal genomes including human as described in ^48^.
- phyloP46way_primate, phyloP (phylogenetic p-values) conservation score based on the multiple alignments of 10 primate genomes including human.

NA values were zero-imputed and each feature was standardized to have mean of zero and variance of one.

##### Response variable

“ClinicalSignificance” was transformed into a binary classification variable with ‘Pathogenic’ corresponding to class 1 and ‘Benign’ corresponding to class 0.

###### Data: dbscSNV

Table S2 from the supplementary material of^49^ was used to train and evaluate models in a 10-fold cross validation (2959 variants, 1164 from the positive class).

**dbscSNV features** (without conservation, described in ^49^)

- ‘PWM_ref, ‘PWM_alt’,
- ‘MES_ref, ‘MES_alt’,
- ‘NNSplice_ref, ‘NNSplice_alt’,
- ‘HSF_ref, ‘HSF_alt’,
‘GeneSplicer_ref, ‘GeneSplicer_alt’,
- ‘GENSCAN_ref, ‘GENSCAN_alt’,
‘NetGene2_ref, ‘NetGene2_alt’,
‘SplicePredictor_ref, ‘SplicePredictor_alt

Kipoi model, conservation and SPIDEX features were the same as for the ClinVar dataset.

##### Response variable

“Group” variable in the original table - ‘Positive’==1 and ‘Negative’==0.

### Meta-model and evaluation

Logistic regression implemented in scikit-learn with default parameters was used to build the meta model using different feature subsets. 10-fold cross-validation was used (implemented in ‘sklearn.model_selection.cross_validate’) to evaluate models using the auROC metric.

### Code availability

Kipoi, kipoi_veff, and kipoi_interpret are available as python packages on PyPI and their source code is available at https://github.com/kipoi/kipoi. https://github.com/kipoi/kipoi-veff and https://github.com/kipoi/kipoi-interpret correspondingly. Models are hosted at https://github.com/kipoi/models. Code to reproduce the results is available at https://github.com/kipoi/manuscript.

## Author contributions

ZA, RK, JI, AS, AK, OS, JG conceived the Kipoi API. ZA, RK implemented the Kipoi API. ZA, RK conceived and implemented kipoi-veff. ZA, RK and AS conceived and implemented kipoi-interpret. ZA, RK, JI, NX, AB performed the analysis. DK compiled the DNA accessibility dataset. ZA, RK, JI, NX, AS, LU contributed models to the repository. AK, OS, and JG designed and supervised research. ZA, RK, AK, OS, JG wrote the manuscript.

## Acknowledgement

We thank Chuan-Sheng Foo for early discussion on the manuscript and Nejc Zupan for implementing the website. We thank Daniel Quang for providing help with FactorNet and trained models for CEBPB and MAFK. We thank Wolfgang Huber for feedback on the manuscript. Z.A. and J.C were supported by a Deutsche Forschungsgemeinschaft fellowship through the Graduate School of Quantitative Biosciences Munich. J.C. was supported by the Competence Network for Technical, Scientific High Performance Computing in Bavaria KONWIHR. L.U. received support from core funding of the European Molecular Biology Laboratory and the European Union’s Horizon 2020 research and innovation programme (grant agreement number N635290). J.I. is supported by a Stanford BioX Fellowship. A.S. is supported by an HHMI International Student Research Fellowship and a Stanford BioX Fellowship. D.S.K. is supported by a Stanford BioX Fellowship. A.B. is supported by NIH grant 1DP2OD022870. A.K. is supported by NIH grants 1DP2OD022870 and 1U01HG009431. This work was supported by NVIDIA hardware grant providing a Titan X GPU card.

